# On the coupling between membrane bending and stretching in lipid vesicles

**DOI:** 10.1101/2024.09.13.612881

**Authors:** Håkan Wennerström, Emma Sparr, Joakim Stenhammar

## Abstract

The formation of a lipid vesicle from a lamellar phase involves a cost in bending energy of 100–1000 times the thermal energy for values of the membrane bending rigidity *κ* typical for phospholipid bilayers. The bending rigidity of a bilayer is however a strongly decreasing function of its thickness *h*, and the bilayer can thus reduce its bending energy by stretching (and thus thinning) the bilayer. In this paper, we construct a simple model to describe this mechanism for the coupling between bending and stretching and analyse its effect on the bending energy and thermal fluctuations of spherical lipid vesicles. We show that the bilayer thinning becomes significant for small vesicles, and for a vesicle with radius *R*_0_ ∼ 15 nm there is a sizeable thinning of the bilayer compared to the planar state. We furthermore demonstrate how this thinning is associated with a significant decrease in free energy due to the thermally excited bending modes. We argue that this previously unexplored effect can explain the experimentally observed lower limit of achievable vesicle sizes, which eventually become unstable due to the thinning of the bilayer. We also sketch how this effect provides a potential generic mechanism for the strong curvature dependence of protein adsorption to lipid membranes.

## 1. Introduction

Biological membranes constitute the basic protective envelope through which cells and organelles interact with their surroundings. Phospholipid membranes are dynamic structures that respond to external stimuli by, for example, changing their shape and adsorbing or releasing small vesicles through fusion or fission [1]. One key characteristic property of the membrane is its local curvature, ℋ. While the undeformed plasma membrane is nearly flat, corresponding to ℋ = 0, many examples of highly curved membranes are found in cellular vesicles and organelles, such as endosomes, exosomes, mitochondrial inner membranes, the tubular network of the endoplasmic reticulum, and in the Golgi apparatus [2]. Membrane structures with very high local curvatures are furthermore common in membrane remodelling processes such as membrane budding and vesicle fusion and fission [3]. These strong local variations of ℋ are not only essential for describing the local structure and deformation the membrane, but also affects its biochemical function. There are several examples of proteins whose association to the membrane can be tuned by local variations in membrane curvature [4, 5, 6, 7, 8], with a key example being amphipathic helices associated with the fusion and fission of membrane vesicles. Down the line, such curvature-dependent protein-membrane interactions also play an important role in controlling the sorting, trafficking and function of membrane associated proteins [4, 5, 6].

In experimental studies of lipid membranes, it is common to use spherical vesicles prepared from one or several lipid components. Small unilamellar vesicles (SUVs) typically have radii between 10 and 50 nm, while giant unilamellar vesicles (GUVs) are in the micrometer size range. Thus, GUVs fall within a similar size range as most cells, and SUVs have a similar size as vesicles used for intra- and extracellular communication. There are several methods to prepare unilamellar vesicles, whose protocols differ depending on the type of vesicles desired. These include sonication and extrusion for the preparation of SUVs [9, 10], and electroformation, gel-assisted swelling and microfluidics-based methods for the preparation of GUVs [11, 12, 13].

Our description of a bent membrane starts from Helfrich’s expression for the bending energy of an elastic membrane [14]

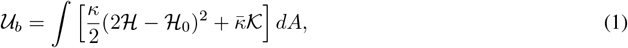

where ℋ is the mean curvature, ℋ_0_ the spontaneous curvature, *κ* the bending rigidity, 𝒦 the Gaussian curvature, and 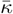 the saddle-splay rigidity. Although Eq. (1) is most frequently used to describe lipid bilayers, it is equally useful when treating lipid and surfactant monolayers. Unlike ℋ_0_ for the bilayer, the monolayer spontaneous curvature ℋ_0_ is generally nonzero even for membranes made from a single lipid component, and is determined by the balance between the head-group repulsion and the volume constraints set by alkyl chain packing. _0_ has also been shown to be a central parameter in the description of the phase behaviour of microemulsion systems [15]. In its basic version, the Helfrich free energy treats the three parameters *κ*, 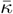, and ℋ_0_ as generic material constants, without any explicit coupling to the molecular details of the membrane, although the same expression can also be derived from a formal thermodynamic description, relating the material parameters to the microscopic stress profiles measured across the mono-or bilayer [16].

For a perfectly spherical, unilamellar vesicle in the absence of spontaneous curvature, corresponding to a symmetric bilayer composed of a single molecular component, the bending energy reduces to the simple expression

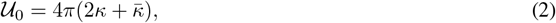

which is independent of the vesicle radius *R*_0_. For *κ* = 30*k*_*B*_*T* and 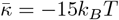, which are reasonable estimates for lipid bilayers [17, 18], the energy 𝒰_0_ ≈ 600*k*_*B*_*T* required for forming a vesicle from an infinite planar bilayer is thus sizeable compared to the thermal energy. However, since a vesicle is a mesoscopic object, the energy is nevertheless a small quantity for GUVs when viewed as an energy per lipid molecule. However, for SUVs near the lower limit of achievable sizes, even the energy per lipid can become sizeable: A vesicle with *R*_0_ = 12 nm and head-group area of 0.7 nm^2^ contains approximately 5200 lipid molecules, implying a bending energy of ∼ 0.1*k*_*B*_*T* per lipid. This constitutes a significant energetic penalty imposed by the vesicle curvature, and there is thus an energetic driving force for releasing some of this frustration, for example by stretching the bilayer.

In continuum treatments of bilayer bending and deformation, the bilayer area *A* is usually assumed to be constant and determined only by the number of lipids in the bilayer. Mathematically, the constant-area constraint is typically accounted for via the use of a Lagrange multiplier in the free energy [19, 20, 21, 22], which can then be viewed as the tension created by a stretching or contraction of the bilayer. In a recent paper [23], we introduced an alternative, statistical-mechanical approach for fulfilling the constant-area condition in GUVs by explicitly imposing that each configuration in the ensemble of vesicle configurations has a predefined area. In the present paper we go beyond the constant-area constraint and highlight a previously unexplored mechanism leading to a coupling between bilayer bending and stretching: Due to the incompressible nature of the bilayer, a stretching of the bilayer leading to an area *A > A*_0_ is associated with a decrease in bilayer thickness *h*, which in turn decreases its bending rigidity *κ*. Using a continuum elastic model for the bending and stretching of a thin sheet, we explore the thermodynamic consequences of this coupling, and show that it will effectively put a lower bound on the achievable size of spherical vesicles. In addition, we discuss how this curvature-dependent effect can provide a basic mechanism for curvature sensing in biomolecules adsorbing on lipid membranes.

## 2. Theory

We first consider a near-spherical, unilamellar vesicle with constant area 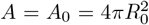, where *R*_0_ is the area of the unperturbed sphere. For each vesicle configuration, the deviation from spherical shape can be characterized by an expansion of the radial interface position *r*(*θ, ϕ*) in spherical harmonics *Y*_𝓁*m*_ as [19]

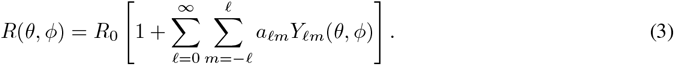

The coefficient *a*_00_ ≤ 0 gives the effective radius while *a*_1*m*_ corresponds to a translation of the vesicle, and the bending energy in Eq. 1 is thus only affected by the expansion coefficients with 𝓁 ≥ 2, according to [19]

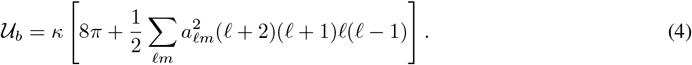

Here, ∑_𝓁*m*_ implies summation over all 𝓁 ≥ 2 and the corresponding *m*. As long as the magnitude of the coefficients *a*_𝓁*m*_ decreases slowly enough with 𝓁, such as when their values are determined by the equipartition theorem [23], 𝒰_*b*_ diverges and it thus becomes necessary to introduce an upper cutoff 𝓁_max_ in Eq. 4, as we discuss further below.

The elastic energy 𝒰_*s*_ associated with small membrane expansions or contractions compared to the optimal area *A*_0_ is given by [24, 25, 26]

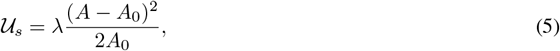

where *λ* is the area expansion modulus, which is typically in the range 0.1 to 0.3 J m^−2^ for lipid bilayers [27, 28, 29]. To leading order in the expansion coefficients *a*_𝓁*m*_, the vesicle area is expressed as [23]

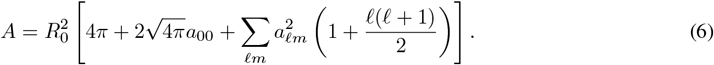

It follows from the final term in Eq. 6 that the area will increase when the vesicle deforms from spherical shape. However, if *a*_00_ *<* 0, corresponding to a decrease in the average radius, the second and third terms can cancel, leading to a deformation without any lateral stretching or contraction of the bilayer. This process is the conventional assumption when describing vesicle fluctuations and deformations, and the resulting coupling between bending and average radius has the consequence that, in the presence of thermally fluctuating bending modes, the vesicle volume *V* shrinks somewhat compared to its value for the unperturbed sphere [23].

According to Eq. 4, the bending energy, and thus the thermal shape fluctuations, for a given mode 𝓁, *m* are independent of *a*_00_, and thus of the vesicle size. Area changes due to stretching or contraction of the bilayer, induced by a change in the value of *a*_00_ in Eq. 6, are thus energetically decoupled from bilayer deformations due to bending. At this level of description, there is therefore no direct coupling between bending and stretching, which we instead introduce in the following through an explicit area dependence of the bending and saddle-splay moduli, *i*.*e*., *κ* = *κ*(*A*) and 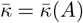.

### Area dependence of the bending modulus

A standard result from the theory of elasticity is that the bending rigidity of a homogeneous elastic sheet of thickness *h* is proportional to *h*^3^ for small deformations [24]. When applied to a membrane, this scaling arises separately for each monolayer if one views each lipid as a linear object interacting with its nearest neighbours at a separation which is shifted from its optimal value due to the bilayer deformation, as shown in Appendix A. Since the two monolayers can slide freely relative to each other, the bending rigidity of the bilayer is simply twice that of the monolayer [25], and it thus follows that *κ* ∼ *h*^3^ also for the bilayer. If the bending rigidity of the monolayer is instead viewed as a surface effect, where the increased energy is a consequence of increasing and decreasing the respective effective areas of the two membrane interfaces, the scaling of instead becomes quadratic in *h*. This has previously been argued to be more accurate for lipid bilayers in the fluid state [29, 27], although quantitative measures of the thickness dependence of *κ* are challenging experimentally. The true exponent is likely system dependent and lies somewhere in between these two extremes, and for the time being we will therefore treat the exponent as an unknown parameter and set *κ* ∼ *h*^*α*^ with 2 ≤ *α* ≤ 3. We will furthermore assume that the bilayer is incompressible, so that the bilayer volume 𝒱_0_ = *hA* remains fixed. This approximation is common in the literature [30, 31, 26] and we expect it to be very accurate since the volume of each hydrocarbon chain remains largely constant far away from any phase transitions. This allows us to write

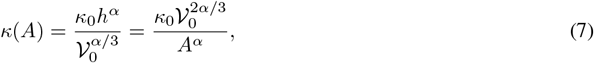

where *κ*_0_ is a generic, *h*-independent constant. Thus, a stretching of the bilayer will lower its bending rigidity, providing a mechanism towards lowering the bending energy. As we show in Appendix B, this coupling term between bending and stretching constitutes the dominant third-order term in a formal series expansion of the free energy about the planar bilayer.

The thickness dependence of the saddle-splay modulus 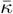 is less straightforward to assess than that of *κ*, and literature estimates for the ratio 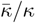 for lipid mono- and bilayers vary with composition [32, 33]. As we show in Appendix C, for a homogeneous, isotropic, and incompressible elastic material, *κ* and 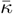 are related as 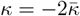, implying that 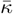 is negative. This relationship is not generically valid for lipid membranes, which are in general both fluid and anisotropic. However, for monolayers whose spontaneous curvature is close to zero, such as for phosphatidyl choline (PC) lipids with more than 14 carbon atoms [34, 35], we argue that this description is reasonable. To proceed, we will therefore make the simple assumption that 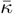 and *κ* are related in the same way as for an isotropic elastic material, so that

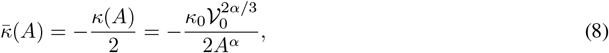

which we thus expect to be a valid approximation for most commonly studied PC vesicles. Finally, the area expansion modulus *λ* is known to vary linearly with *h* for a homogeneous and isotropic medium; however, since this area dependence is significantly weaker than that of *κ* and 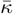, including an explicit area dependence of *λ* leads to higher-order effects that we ignore for simplicity.

The thickness dependence of the saddle-splay modulus 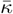 is less straightforward to assess than that of *κ*, and literature estimates for the ratio 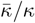 for lipid mono- and bilayers vary with composition [32, 33]. For a homogeneous, isotropic, and incompressible elastic material, *κ* and 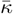 are related as 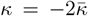 (see Appendix C), implying that 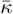 is negative. We argue that this relationship between *κ* and 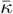 is relevant for monolayers with spontaneous curvatures close to zero, such as those composed of phosphatidyl choline (PC) lipids with more than 14 carbon atoms [34, 35]. Just as for the bending modulus, 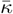 for a symmetric bilayer is twice its value for the corresponding monolayer, since the two leaflets can slide freely with respect to each other. To proceed, we will therefore make the simple assumption that 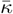 and *κ* for the bilayer membrane are related in the same way as for an isotropic elastic material, so that

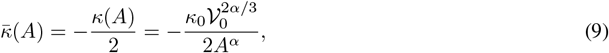

which we expect to be a valid approximation for many commonly studied PC bilayers. In contrast, we expect it to be less useful for membranes composed of lipids with shorter chains, where ℋ_0_ *>* 0, or lipids with highly unsaturated chains, for which ℋ_0_ *<* 0 [34, 35]. In either case, since (*i*) the bending and Gaussian curvature terms are additive in Eq. (2), (*ii*) 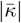 is typically smaller than *κ* and (*iii*) we do not expect 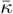 to have a stronger *h* dependence than *κ*, the choice of scaling relation made for 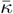 should only have a marginal effect on our results and not affect our qualitative conclusions. Finally, the area expansion modulus *λ* is known to vary linearly with *h* for a homogeneous and isotropic material [19]; however, since this area dependence is significantly weaker than that of *κ* and 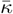, the inclusion of an explicit area dependence of *λ* leads to higher-order effects that we ignore for simplicity.

From Eqs. 2, 5, and 9 the total bend-stretch energy for forming a spherical vesicle of area *A* is

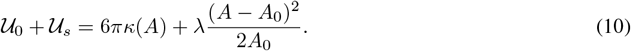

The optimal vesicle area is obtained by inserting Eq. 7 into Eq. 10 and minimising with respect to *A*, yielding

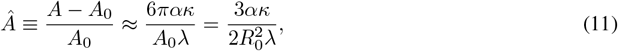

where we have assumed that *A* ≈ *A*_0_ in the denominator. Equation 11 can be further simplified by using the approximate relation [25] *κ* = *λh*^2^*/*12 between the bending and stretching moduli, leading to

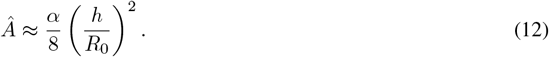

Notably, the scaling exponent *α* only enters the area expansion as a linear factor multiplying *Â* and does not affect its scaling with *R*_0_ or *h*. The results are therefore only moderately shifted as *α* varies between 2 and 3, and in the following we therefore set *α* = 3 for simplicity. In Fig. 1, we show the relative area expansion as a function of *R*_0_ for three different values of *κ* plotted using Eq. 11, together with the simplified expression in 12; note that the relative decrease in bilayer thickness *ĥ* can be directly obtained from the area expansion as *ĥ* = −*Â*. These results clearly show that, for SUVs approaching *R*_0_ = 15 nm, we do expect bilayer expansions in excess of one percent, which is potentially high enough to cause vesicle rupture [26].

**Figure 1:**
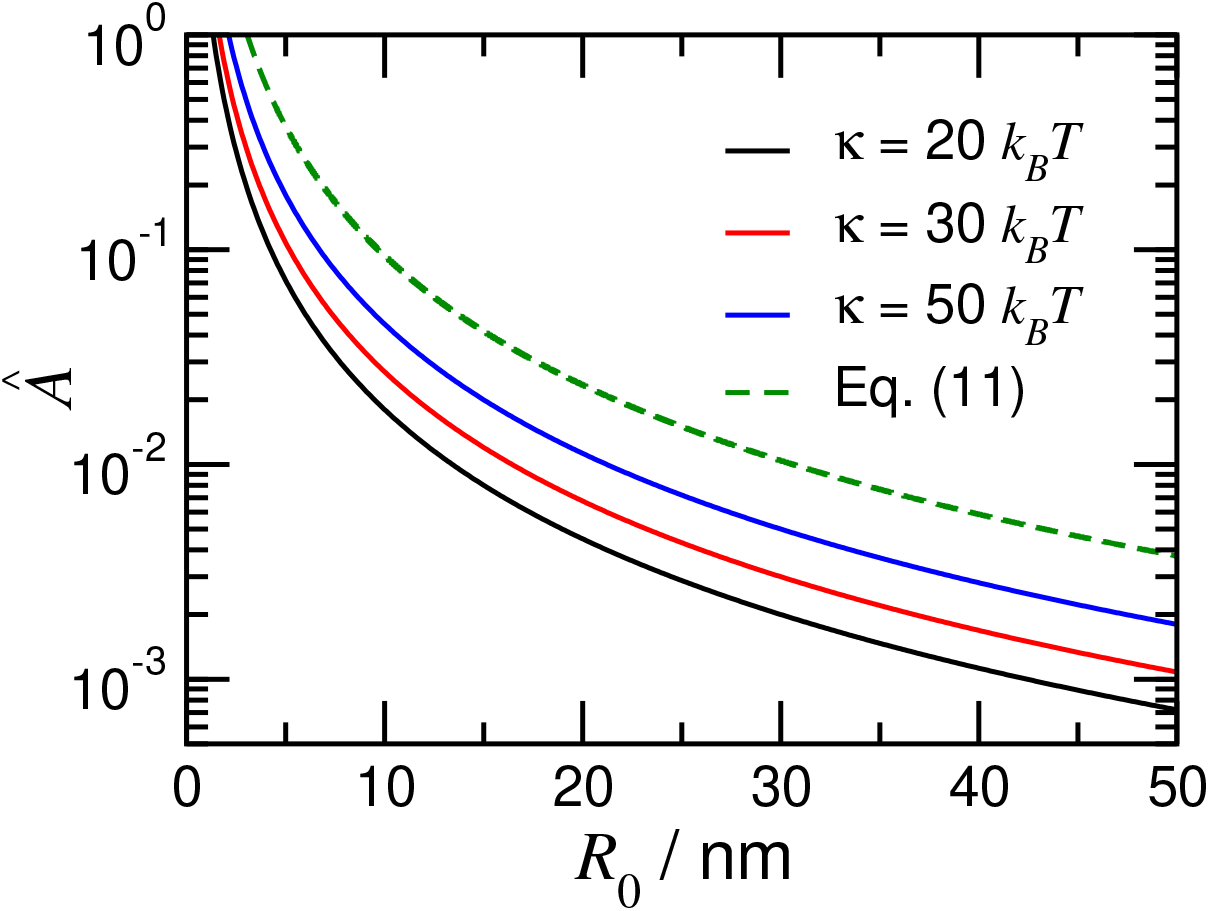
Relative vesicle area increase *Â* as a function of the vesicle radius *R*_0_. The solid curves were obtained from Eq. 11 the curves using *α* = 3, *λ* = 0.2 J m^−2^ = 50*k*_*B*_*T/*nm^2^ and indicated values of *κ*. The dashed line was obtained from Eq. 12 with *h* = 5 nm.

### Consequences for thermal shape fluctuations

The analysis leading to Eq. 11 is based on the assumption of a perfectly spherical vesicle, thus ignoring shape fluctuations due to thermally excited bending modes. Since the amplitude of these fluctuations depend on the bending energy of each mode, the coupling between bending and stretching will also affect the fluctuation spectrum [20, 21]. From Eq. 4 and the equipartition theorem, in the absence of any coupling between bending and stretching each fluctuating bending mode (𝓁, *m*) is associated with an average bending energy

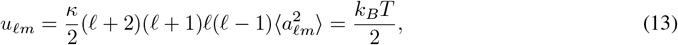

with 𝒰_*b*_ = ∑_𝓁*m*_ *u*_𝓁*m*_ and ⟨ … ⟩ denotes a statistical-mechanical average. The bending energy can in turn be associated with a free energy ℱ_*b*_ = −*k*_*B*_*T* ln *Z*, with the configuration integral *Z* given by

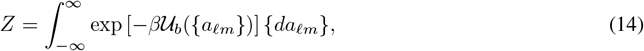

where *β* = (*k*_*B*_*T*)^−1^ is the inverse thermal energy. Note that, unlike in our previous study [23], Eq. 14 puts no constraint on the total volume *V* of the vesicle, which we previously introduced *via* a *δ* function in the configuration integral in order to study the effect of external osmotic gradients on the bending fluctuations [23]. Thus, Eq. 14 is approximate in that it decouples the different fluctuation modes, which makes the integral factorisable. This relaxation of the constant-volume condition leads to a slight overestimation of the magnitude of ℱ_*b*_, although we do not expect it to affect the results radically. Using Eq. 4, the product of Gaussian integrals can be solved to give

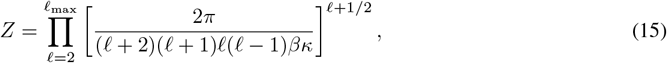

where we have used the fact that, for each 𝓁, there are 2𝓁 + 1 degenerate fluctuation modes. The corresponding free energy is given by

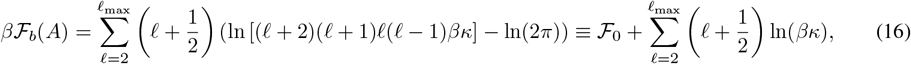

where ℱ_0_ contains all the terms that are independent of *κ* (and thus of *A*). Using Eq. 7, the free energy change Δℱ_*b*_ associated with an area change from *A*_0_ to *A* becomes

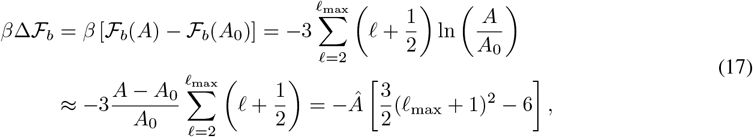

where, in the third equality, we have assumed small enough deviations from the optimal area *A*_0_ to enable linearisation of the logarithm. In contrast to the direct stretching energy of Eq. 5, this contribution is linear in (*A* − *A*_0_), and promotes an expansion of the bilayer due to the resulting decrease in bending energy. The precise value ofΔℱ_*b*_ is strongly dependent on the choice of cutoff 𝓁_max_, which should be selected as the mode where the continuum description of deformations is no longer valid. For an SUV with *R*_0_ = 12 nm and *h* = 5 nm, using 𝓁_max_ ∼ *R*_0_*/h*, which we previously used to estimate 𝓁_max_ for GUVs [23], yields 𝓁_max_ = 2, which is unphysically small. Instead using 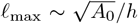 yields 𝓁_max_ ∼ 9, while an estimate of 𝓁_max_ from the number of *molecular* degrees of freedom gives 𝓁_max_ values about an order of magnitude higher, which is clearly an overestimation. Due to this broad range of relevant 𝓁_max_ values, in Fig. 2, we plot *β*Δℱ_*b*_ as a function of 𝓁_max_, calculated from Eq. 17 for four realistic values of the area expansion *Â*. Clearly, the free energy gain from expanding the vesicle area can be significantly larger than the thermal energy for realistic values of 𝓁_max_, indicating that, for SUVs near their lower size limit, there is a significant driving force for thinning the bilayer enough to cause vesicle rupture. In the following, we will discuss this and other potential experimental consequences of the coupling between bending and stretching.

**Figure 2:**
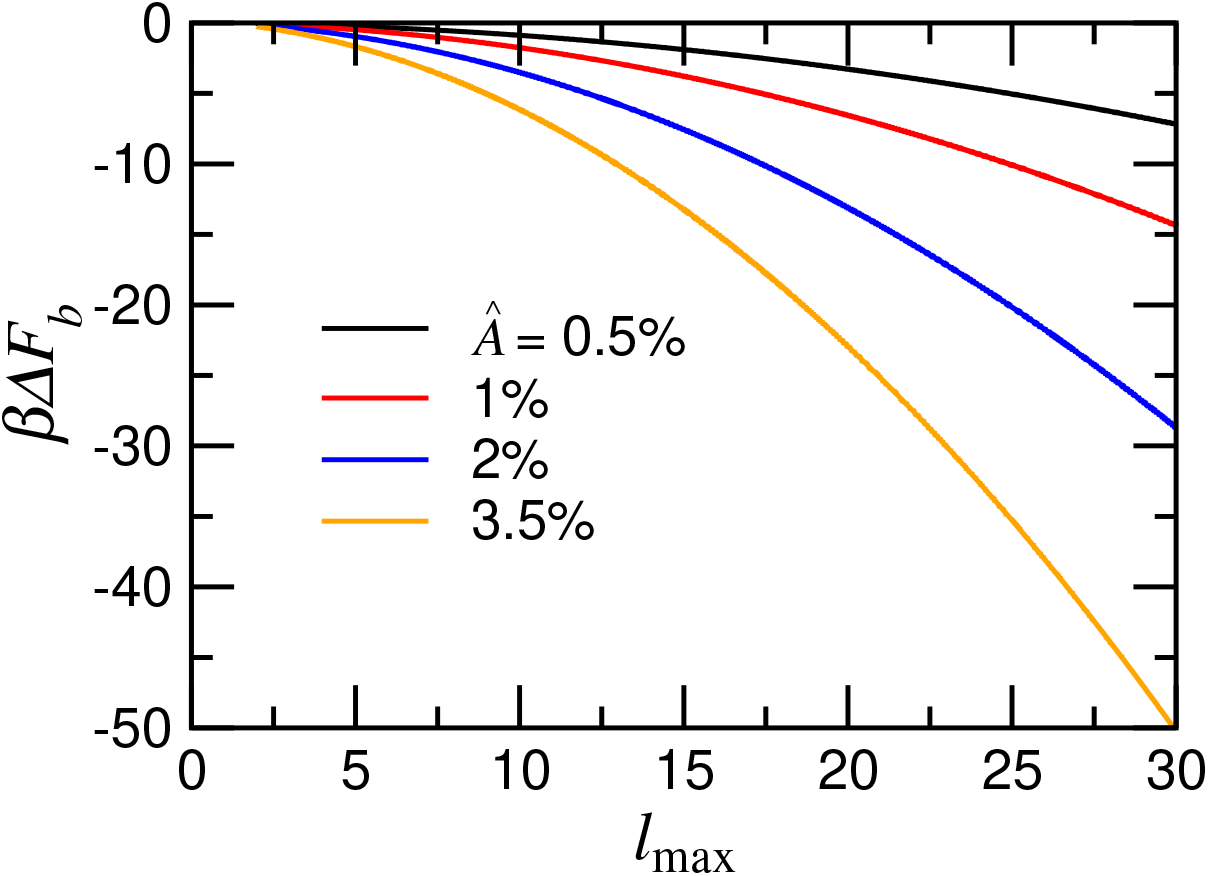
Free energy change *β*Δℱ_*b*_ from increased thermal fluctuations due to vesicle stretching, as given by Eq. 17 for various values of the area expansion *Â* (*c*.*f*. Fig. 1). Note the strong dependence on the cutoff parameter 𝓁_max_.

## 3. Discussion

The main aim of this study is to develop a simple model describing the energetic coupling between membrane bending and stretching. The derived relations are based on the key assumption that the bending rigidity *κ* is a decreasing function of the bilayer thickness *h*, which is consistent with experimental observations [27], albeit with uncertainties in the precise mathematical relationship. While the model is clearly idealised, our results show that such a generic coupling has significant energetic consequences for small vesicles in the 10 − 20 nm range, since already a 2 percent expansion of the bilayer is associated with a free energy reduction significantly larger than *k*_*B*_*T* for reasonable values of the upper cutoff 𝓁_max_. From an experimental perspective, a decrease in bilayer thickness of a few percent is small and therefore difficult to demonstrate directly with sufficient accuracy. Instead of directly measuring the bilayer thickness one can make an estimate through some property that depends strongly on the membrane thickness. One such potential property is the order parameter *S* of the hydrocarbon chains, which is known from simulations to vary strongly with *h* [36]. This parameter can readily be estimated from NMR relaxation experiments, since the *T*_2_ relaxation time of ^1^H, ^2^H and ^13^C nuclei in the hydrocarbon tails has a quadratic dependence on *S* [37, 38]. Some experimental studies have indeed measured a significant difference of the *T*_2_ values between planar bilayers and SUVs, although the origin of this difference is disputed [39]. Thus, further careful experiments are necessary to establish this potential connection between curvature, bilayer thickness, and chain ordering. In the following, we will finish by discussing two potential further consequences of the vesicle thinning effect, namely (*i*) the experimentally observed lower size limit for vesicle stability, and (*ii*) the curvature-dependent binding of proteins to lipid bilayers.

### Lower limit of vesicle stability

Experimentally, the size distribution of vesicles has been observed to sensitively depend on the preparation procedure, and even differ depending on details in the experimental protocol for the same lipid composition and preparation method [11, 12, 13]. This shows that the observed vesicle size distribution is in fact not an equilibrium property, which would be determined purely by the vesicle free energy, but rather represents a metastable state where vesicle fusion and Ostwald ripening is slow. In practice it is difficult to make vesicles with a radius smaller than 10 nm regardless of the preparation method [40, 41, 42, 43], indicating that, unlike the overall size distribution, the minimal vesicle size is in fact determined by some general mechanism. It has previously been suggested that this lower size limit is determined by a fourth-order curvature energy term [44, 25]. Our results however provide a potential explanation for this lower stability limit of SUVs: since the bending rigidity *κ* decreases with bilayer thickness, there is a considerable driving force for bilayer stretching, as quantified by Eq. 11. As the bilayer stretches, the area per head-group increases and exposes a larger portion of the hydrophobic interior, lowering the barrier and increasing the driving force for vesicle fusion and eventually causing rupture. If the lower radius limit *R*_*c*_ of SUVs is indeed determined by this mechanism, the stability with respect to fusion will be a continuously increasing function of the vesicle size, and vesicles with radii only slightly above *R*_*c*_ will have a lower barrier for fusion than vesicles with *R* ≫ *R*_*c*_. The kinetic stability of these smaller vesicles will thus be more sensitive to perturbations in the system, such as the presence of impurities or surfaces.

### Curvature sensing

Proteins that preferentially bind to membranes of large (local) curvature are referred as curvature sensing proteins. The ability of proteins to distribute between membrane surfaces based on differences in membrane curvature has previously been associated with various protein functions, such as reaction rates of lipid synthesis, vesicle fission, and protein polymerisation [45, 46, 47]. Although the term “curvature sensing” implicitly infers a mechanism relying on the recognition of a certain membrane geometry or shape, there are several examples where the underlying molecular mechanism rather arises from differences in the lipid packing density in the membrane and/or repulsion between the adsorbed proteins [48, 6, 49, 50, 51, 52]. As we demonstrate in this study, an increase in membrane curvature is expected to yield an increase of the effective area per headgroup. The lateral stretching that occurs for SUVs thus implies a locally increased exposure of hydrophobic hydrocarbon chains, sometimes referred to in the literature as “membrane defects”, and provides a potential mechanism towards enhanced protein binding. In the following, we will attempt to quantify this putative generic curvature sensing mechanism.

The area per headgroup in a phospholipid bilayer is typically determined by a balance between three factors: (*i*)the space needed to accommodate the hydrocarbon chains in the interior region, (*ii*) the headgroup repulsion, and (*iii*) the hydrophobic energy due to contact between water molecules and exposed hydrocarbon chains at the polar-apolar interface [53]. For a flat bilayer, the latter contact area can be estimated from a comparison between the change in the effective area per lipid molecule when going from an ordered gel phase, where the exposed hydrophobic area is minimal, to a positionally disordered, liquid crystalline lamellar phase. As an example, for a liquid disordered bilayer composed of dioleoylphosphatidylcholine (DOPC) the effective area per headgoup is above 70 Å^2^, while the corresponding value for gel phase phosphatidylcholine bilayers is below 50 Å^2^ [54]. From these numbers, we estimate the exposed hydrophobic area in the DOPC bilayer to be approximately 30 percent of the total area. For a strongly curved bilayer, such as in SUVs, we previously showed that the bending energy induces an area expansion of order a few percent. However, since the area of the polar head groups remains unchanged by curvature, this area expansion is primarily manifested as an increase in the hydrophobic contact area. Thus, a relative area expansion of 2 percent leads to an 8 percent increase in the area of exposed hydrocarbon chains.

As an example, we will consider the initial stage of a protein adsorbing onto the unperturbed bilayer surface by forming an *α*-helix, a process typically driven by hydrophobic interactions. From the perspective of the bilayer, the protein binding yields an energetically favourable reduction of the exposed hydrophobic area, which from the above argument will be more significant for small vesicles with expanded areas than for flat, unstretched bilayers. If the initial area expansion is 2 percent, the additional eliminated hydrophobic contact area per lipid due to the adsorption is 0.02*a*_0_ ≈ 1.4 Å^2^, assuming an area per headgroup *a*_0_ of 70 Å^2^. Using a hydrophobic surface energy of 30 × 10^−3^J m^−2^, we thus estimate that a vesicle with a 2 percent area expansion will cause an additional adsorption energy of approximately 0.1*k*_*B*_*T* per lipid. Since the number of lipids involved in the association is at least of order 10, this conservative estimate shows that the shift in adsorption energy can be comparable to *k*_*B*_*T*, and the association constant can therefore increase by at least a factor of 2 solely due to this curvature-induced pre-stretching of the membrane. It should finally be noted that the presence of this generic mechanism does not exclude additional, molecularly specific mechanisms for curvature sensing. One example of such a specific mechanism is the preferential bending of *α*-helices, which leads to long helices preferentially adsorbing onto bent bilayers relative to planar ones, with the adsorption of the helical protein *α*-synuclein on lipid vesicles as an important example [55, 56].

### Summary

In this study, we have demonstrated and discussed a number of putative thermodynamic, kinetic, and biological consequences of a physically motivated coupling between bilayer thickness *h* and bending rigidity *κ*. The experimental consequences of this bilayer thinning clearly need to be confirmed by further careful experiments, but we nevertheless believe that the simple and generic nature of the mechanism makes it plausible as an important factor in the energetics and mechanics of highly curved membrane structures in biological settings as well as in simplified model systems.

## APPENDIX A: Thickness dependence of the bending rigidity for a lipid monolayer

We consider a single, planar monoloayer of thickness *h/*2 and describe each lipid as a linear object interacting with its nearest neighbours with an energy *u* per unit length, as sketched in Fig. 3. For small deformations from the flat configuration, we can describe *u*(*d*) as a harmonic potential:

**Figure 3:**
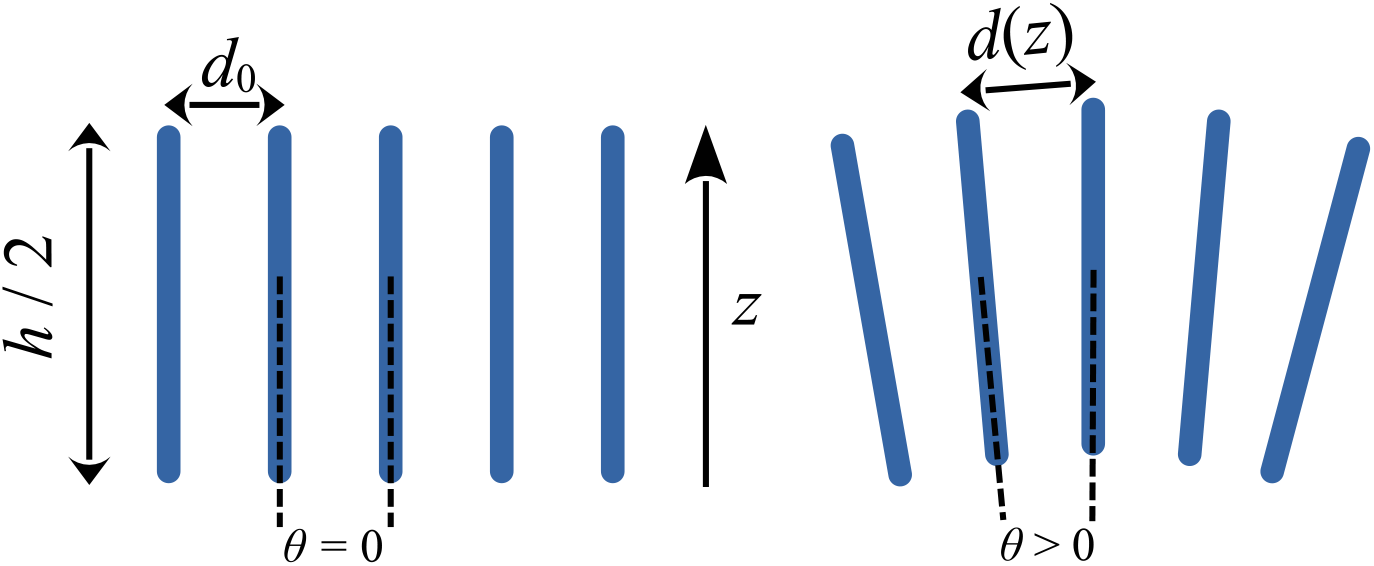
Schematic model used to calculate the thickness dependence of the bending rigidity: Each lipid is described as a linear rod interacting with its nearest neighbours with a harmonic potential, as described by Eq. 19. When the monolayer is bent, each lipid is tilted at an angle *θ* relative to its nearest neighbours, yielding a deformation energy proportional to *h*^3^.

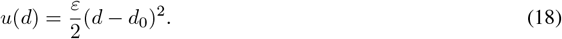

Here, *ε* measures the magnitude of the interaction, *d* is the separation between two points on adjacent lipids, and *d*_0_ their equilibrium separation in the undeformed state. When the monolayer is bent, adjacent lipids become tilted at an angle *θ* with respect to each other, and their total pair interaction energy becomes

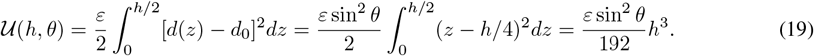

The total energy of the deformed monolayer is given by the sum of all the nearest-neighbour interactions, and thus the bending energy 𝒰_*b*_ of each monolayer also scales as *h*^3^.

## APPENDIX B: Series expansion of the free energy for an elastic membrane

As in Eq. 1, the free energy 𝒰 of the bilayer can be written an area integral over a local free energy density *u*(*x, y*), which we consider a function of the local vertical displacement of the surface *ζ*(*x, y*) and lipid area density *ρ*(*x, y*). We furthermore assume that the bilayer is fully symmetric, corresponding to a single-component lipid composition, leading to ℋ_0_ = 0. The curvature can be characterised by the second derivatives *ζ*_*αβ* ≡_ ∂^2^*ζ/*(∂*α*∂*β*). In the principal coordinate system the cross derivatives vanish, so that we can write

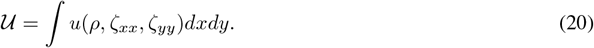

We now use a symmetric, planar bilayer with equilibrium area density *ρ*_0_ as the reference state and expand *u* to second order as

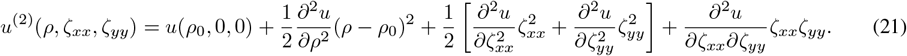

Here, we have used that first-order derivatives with respect to *ζ* vanish by symmetry and ∂*u/*∂*ρ* = 0 since we expand about the equilibrium state. We furthermore have that 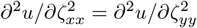 by symmetry. The bending moduli can now be identified from Eq. 1 using the definitions ℋ ≡ (*ζ*_*xx*_ + *ζ*_*yy*_)*/*2 and 𝒦 ≡ *ζ*_*xx*_*ζ*_*yy*_, leading to

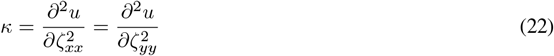

and

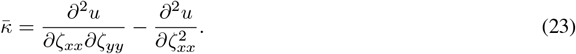

The stretching modulus *λ* can be identified with Eq. 5 if we first assume membrane deformations to be small enough that the area density remains uniform, so that we can replace *ρ*(*x, y*) with its average value *ρ*_av_ = *N/*(2*A*), where *N* is the total number of lipid molecules in the bilayer. For small stretching deformations we can then write (*ρ*_av_ − *ρ*_0_)^2^ ≃ *N* ^2^(*A* − *A*_0_)^2^*/*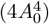, leading to the identification

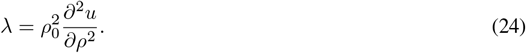

At leading order, there is thus no term that couples between bending and stretching. Such terms arise instead at third order, where we have the additional contributions

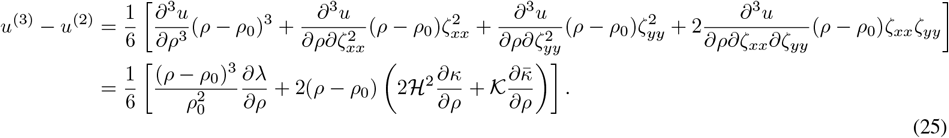

For a membrane with a large resistance towards expansion and compression, the term cubic in (*ρ* − *ρ*_0_) will be negligible, and thus the only sizeable contribution to the bend-stretch coupling at third order is given by the area dependence of *κ* and 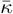, as approximated by Eqs. 7 and 9. It should be noted that we expect Eqs. 21 and 25 to be accurate for spherical vesicles where the local variation in curvature and lipid density is modest. To describe budding and other processes with large variations in local curvature, one should go beyond the uniform density description and explicitly consider the covariation between *ρ*, ℋ, and 𝒦.

## APPENDIX C: Relationship between *κ* and 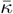 for an isotropic elastic material

In the principal coordinate system, the general expression 𝒰_*b*_ for the bending energy of a deformed elastic plate of thickness *h* is given by [24]

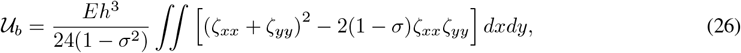

where *σ* is the Poisson ratio and *E* Young’s modulus and we have used the notation of Appendix B. For an incompressible medium, *σ* → 1*/*2 and *E* → 3*µ*, where *µ* is the shear modulus, reducing the number of material constants from two to one. Using the definitions of ℋ and 𝒦 from Appendix B, we can rewrite Eq. 26 in this limit as

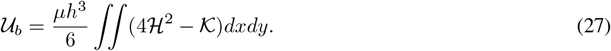

We can thus readily identify the bending modulus *κ* and saddle-splay modulus 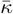 from Eq. 1 as

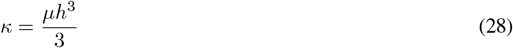

and

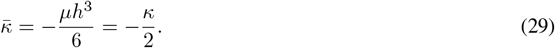

Note that, since Eqs. 28–29 refer to the lipid *monolayer*, the shear modulus *µ* does not vanish, unlike for the bilayer. This is because, even though a lipid monolayer can be considered as a 2-dimensional fluid in the sense that molecules can diffuse laterally, it does not possess the mechanical properties of a simple liquid such as a hydrocarbon, for which the shear modulus is zero. In contrast to the simple liquid, the lipid monolayer cannot be sheared without distorting its molecular structure, and, as a consequence, is able to support shear forces.

## Author Contributions

HW, ES and JS designed the research and co-wrote the paper. HW and JS performed the calculations.

## Acknowledgements

JS and ES kindly acknowledge funding from the Swedish Research Council (grant IDs 2019-03718 and 2019-05296, respectively).

